# Differential virulence contributions of the efflux transporter MexAB-OprM in *Pseudomonas syringae* infecting a variety of host plants

**DOI:** 10.1101/2020.03.04.959429

**Authors:** Tyler C. Helmann, Dana M. King, Steven E. Lindow

## Abstract

Efflux transporters such as MexAB-OprM contribute to bacterial resistance to diverse antimicrobial compounds. Here, we show that MexB contributes to epiphytic and apoplastic growth of *Pseudomonas syringae* strain B728a, as well as lesion formation in common bean (*Phaseolus vulgaris*). While a *mexB* deletion mutant formed fewer lesions after topical application to common bean, these lesions contain the same number of cells (10^5^ to 10^7^ cells) as those caused by the wild-type strain. The internalized population size of both the WT and the MexB mutant within small segments of surface-sterilized asymptomatic portions of the leaves varied from undetectably low to as high as 10^5^ cells/cm^2^. Localized populations of bacteria within the leaf must apparently exceed a threshold size of about 10^5^ cells/cm^2^ in order for a visible lesion to form. Strain B728a was capable of moderate to extensive apoplastic growth in diverse host plants including lima bean (*P. lunatus*), fava bean (*Vicia faba*), pepper (*Capsicum annuum*), *Nicotiana benthamiana*, sunflower (*Helianthus annuus*), and tomato (*Solanum lycopersicum*). MexB was not required for growth in a subset of these plant species, and in which the duration of growth after inoculation was longer, indicating apparent variation in onset time or magnitude of plant chemical defenses among those hosts. The use of a hyper-susceptible efflux pump mutant strain is an informative strategy to explore the diversity of host chemical immune responses.

## Introduction

Plants produce diverse specialized antimicrobial metabolites to protect against potential pathogens and pests (Piasecka et al. 2015). The identity and abundance of these compounds differ between plant clades (Dixon 2001). Together, they constitute a chemical defense that potential plant pathogens must detoxify or tolerate to successfully colonize a host. Most antimicrobial compounds derived from plants can be placed into one of several chemical classes, including phenolics, terpenoids, alkaloids, and lectins/polypeptides (Cowan 1997). These specialized metabolites may be either preformed (“phytoanticipins”) or synthesized *de novo* in response to a pathogen (“phytoalexins”), although this distinction is poorly defined for many compounds (Dixon 2001). Plant-produced antimicrobial metabolites may target many cellular targets including plasma membranes, proteins, or nucleic acids in potential plant pathogens (Piasecka et al. 2015).

Bacteria have evolved several mechanisms to tolerate the presence of plant-produced antimicrobial compounds. These traits can therefore be considered necessary compatibility factors enabling the colonization of a given habitat or plant host. For example, *Pseudomonas syringae* DC3000 requires several genes to overcome the toxicity of aliphatic isothiocyanates produced by *Arabidopsis* and these genes confer resistance to strains that are otherwise incapable of tolerating these molecules (Fan et al. 2011). These “<u>s</u>urvival in *<u>A</u>rabidopsis* e<u>x</u>tracts”, or *sax,* genes encode both detoxifying enzymes as well as components of a transporter in the resistance-nodulation-division (RND) efflux protein family (Fan et al. 2011). Members of the RND family of transporters have been widely studied due to their large contribution to bacterial antibiotic resistance *in vitro.* Only now is there beginning to be further understanding of their broader role in other environmental habitats (Alvarez-Ortega et al. 2013). By far, the best-characterized RND transporters are AcrAB-TolC in *Escherichia coli* and MexAB-OprM in *Pseudomonas* species (Nikaido 1996). Related transporters have been shown to be required for full virulence of several model plant-colonizing bacterial, including *Ralstonia solanacearum*, *Erwinia amylovora*, and *Xylella fastidiosa* (Brown et al. 2007; Burse et al. 2004; Reddy et al. 2007). A common strategy to identify the targets of efflux transporters involves documentation of increased tolerance of various antimicrobial compounds when the pumps are overexpressed (Lomovskaya et al. 1999; Sulavik et al. 2001). While such strategies are an efficient method for identifying potential substrates for efflux pumps, they do not always reveal the biological significance of such exporters. The local repressor of MexAB-OprM (PmeR) in *P. syringae* DC3000 responds to plant secondary products such as flavonoids (Vargas et al. 2011), suggesting that these molecules are true substrates rather than artifacts of non-specific substrate binding sometimes seen when efflux transporters are overexpressed. Members of the RND protein family are responsible for the removal of a diverse range of toxic substrates from the cytoplasm of Gram-negative bacteria. However, relatively little is known about the breadth of compounds that they are able to transport, particularly for the large number of naturally occurring inhibitory compounds. Because of this, there is a lack of insight as to whether these transporters function in an environmentally-specific manner. In this study we use a hyper-susceptible *mexB* mutant of *Pseudomonas syringae* strain B728a as a bioreporter to examine differences in chemical defense operative against a plant pathogen capable of colonizing several susceptible host plant species.

*P. syringae* pv. *syringae* B728a (B728a) is pathogenic to common bean (*Phaseolus vulgaris*) and *Nicotiana benthamiana* (Vinatzer et al. 2006), and was recently reported to cause disease symptoms on pepper (*Capsicum annuum*) (Morris et al. 2019). Strain B728a has also been reported to be a poor colonizer of tomato (*Solanum lycopersicum*) (Lin and Martin 2007). In strain, B728a MexAB-OprM (*Psyr_4007-9*) is required for full virulence on common bean (Stoitsova et al. 2008) and homologs in *P. syringae* strains DC3000 and 1448A conferred resistance to a range of toxic compounds including antibiotics, plant-derived antimicrobial compounds, detergents, and dyes, presumably by exporting them from the cell (Stoitsova et al. 2008). MexB is located in the inner membrane, and is responsible for substrate specificity of the protein complex (Tikhonova et al. 2002), while the outer membrane protein OprM functions also with other RND efflux systems to enable passage of toxicants from the cytoplasm to the exterior of the cell (Schweizer 2003). The process of infection of plants by foliar phytopathogens such as *P. syringae* has received considerable attention. Typically, an initial epiphytic colonization stage facilitates sufficient multiplication of immigrant bacteria on leaves to enable subsequent invasion of the leaf interior, that can be followed by growth in the apoplast, ultimately leading to disease symptoms on leaves under favorable environmental conditions and if plants are susceptible to infection (Xin et al. 2018). Foliar lesions result from the localized multiplication of bacteria inside leaves. The release of water from plant cells into intercellular spaces in leaves creates watersoaking that facilitates release and distribution of nutrients from plant cells, is apparently induced by effectors produced by bacteria (Xin et al. 2016). The assembly of large numbers of bacteria within leaves often causes chlorosis and subsequent necrosis and dehydration in the lesions (Leben 1981). Bacteria within disease lesions can function as the source of inoculum for spread of secondary inoculum via water splash or aerial dispersal (Arnold et al. 2011). While there has been extensive study of bacterial traits needed for growth within plants after their introduction into the plant interior, there has been relatively little study of the early stages of infection, particularly the transition between epiphytic and endophytic populations that lead to disease symptoms. The survival of bacterial cells in lesions as they age, a site where inhibitory compounds might be expected to be found has also not received much attention. Here we examine these various stages of bacterial growth and survival *in planta* that are epidemiologically important, with special attention to the role of efflux transporters such as MexAB-OprM. Most previous attention to the process of infection has focused on particular model plants that interact with a given pathogen. Many plant pathogenic bacteria, particularly those in *P. syringae* phylogroup 2, are now recognized to have relatively broad host ranges (Morris et al. 2019), and yet it remains unclear to what extent a given virulence factor such as an efflux transporter would contribute to virulence on plants that would be expected to differ substantially in their chemical defenses. In this study, we will address the extent to which a given *P. syringae* strain is a promiscuous colonizer of a variety of plants and the relative importance of MexAB-OprM to its virulence on these hosts.

Within the genus *Phaseolus*, five species have been domesticated. There is also abundant diversity within common bean (*P. vulgaris*) since it was domesticated from two geographically divergent gene pools (Kwak and Gepts 2009). The range of susceptibility within *Phaseolus* species to infection by compatible bacterial pathogens indicates a level of quantitative resistance, that might readily be explained by variation in their content of antimicrobial secondary metabolites (Poland et al. 2009). We hypothesized that due to this variation, not all antimicrobial metabolites present in a given plant genotype would be substrates of MexAB-OprM and that differential MexB-dependent growth of strain B728a would be apparent among such plants. By examining the growth of the hyper-susceptible B728a Δ*mexB* mutant in a variety of plant species, we addressed the apparent differential role of chemical defenses in these important crop plant species.

## Results

### MexB contributes to fitness on the leaf surface

Given that B728a Δ*mexB* had been shown to be hyper-susceptible to diverse antimicrobial compounds (Stoitsova et al. 2008), we reasoned it might be an effective bioreporter strain to detect chemical toxicants present in and on the leaves of various plant species. The MexAB-OprM efflux pump is apparently completely dispensable for growth in the rich medium King’s B (KB) since no difference in growth was seen between it and the wild type strain (Fig. S1). However, when sprayed onto the leaves of the susceptible common bean variety Bush Blue Lake 274, subsequent epiphytic bacterial populations established by Δ*mexB* after two days incubation on moist leaves were significantly lower than that of the wild-type (WT) strain (Fig. 1). This suggests that an inhibitory compound was present on the leaf surface, necessitating the role of the MexAB-OprM efflux transporter for full fitness in this habitat.

**Figure 1.**
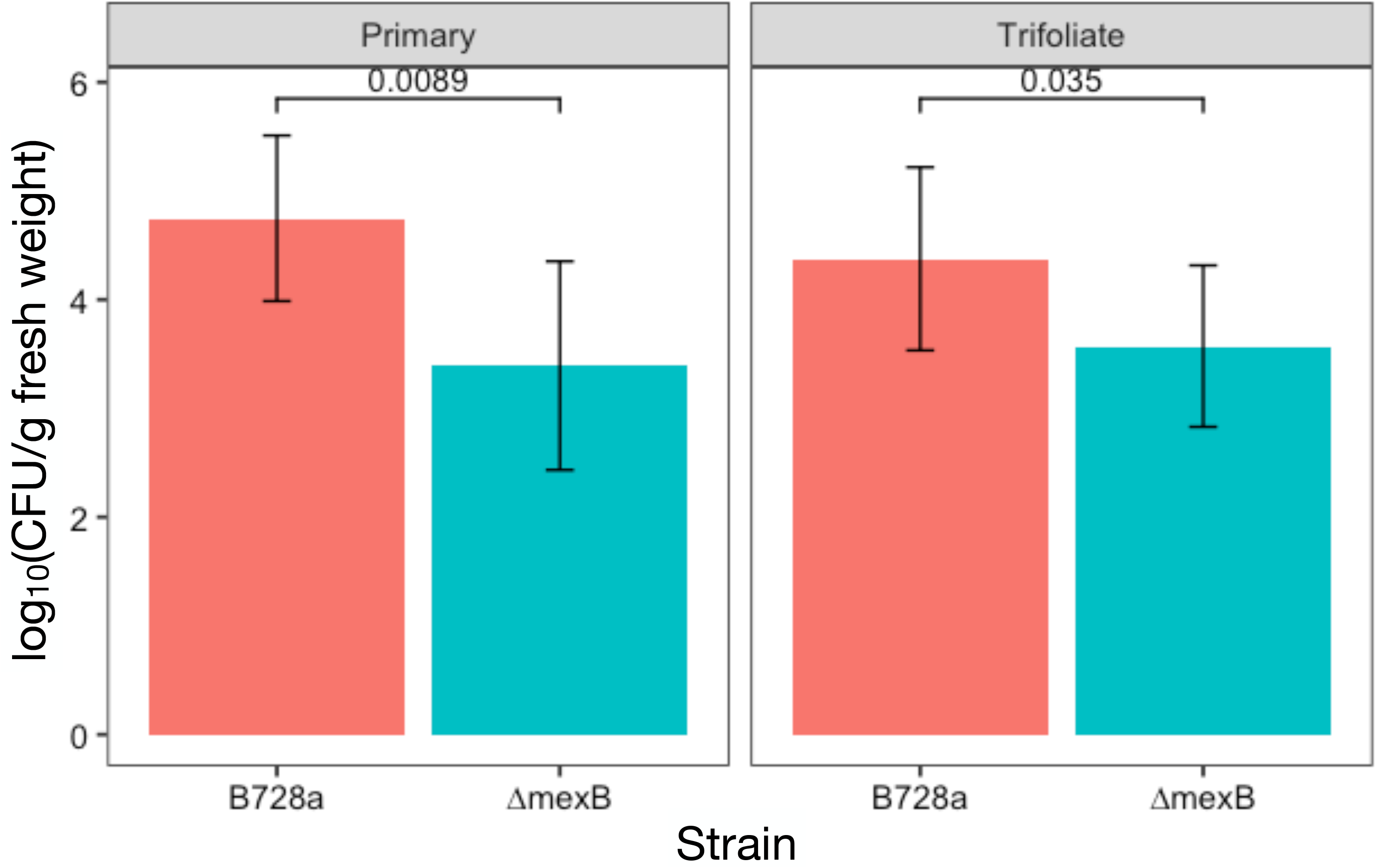
B728a Δ*mexB* establishes lower population sizes on the surface of common bean leaves than the WT strain. For the two leaf types tested, epiphytic populations at high humidity were significantly different two days following a spray inoculation (Wilcoxon signed-rank test). Mean CFU per g leaf tissue are shown, with error bars indicating the standard deviation. N = 10 per strain per leaf type.

### MexB contributes to late-stage multiplication and survival in common bean

When inoculated directly into the apoplast of common bean, the population size of both B728a WT and Δ*mexB* increased similarly with time for two days (Fig. 2). While little additional increase in population size of the WT strain occurred after two days, the population size of Δ*mexB* tended to decrease with time and its population in the apoplast at 6 and 8 dpi was significantly lower than that of wild type strain (Fig. 2). The plateauing of population sizes after two days incubation, particularly for that of the WT strain suggested either that this maximal apoplastic population reflected an intrinsic carrying capacity of the leaf interior for this species or the intervention of an active immune response from the plant that limited further growth after such a time. To discriminate these scenarios we inoculated the interior of leaves with bacterial inoculum of varying concentrations and measured the resultant bacterial population sizes after four days; sufficient time for even low initial numbers of bacteria to reach high internal leaf populations. Importantly, irrespective of the number of cells initially introduced into the leaf interior, bacterial population sizes increased approximately 1000-fold and thus were highly correlated with the original inoculum concentration (Pearson’s correlation coefficient = 0.90) (Fig. S2). Such a pattern of growth is inconsistent with the concept of a resource-limiting carrying capacity and instead is highly supportive of a model in which, irrespective of their initial concentration in the plant, bacterial cells are able to grow only for a limited period of time (ca. 2 days) before growth is halted. Such a pattern likely reflects a delayed but effective resistance response in the host, and because of hypersensitivity of Δ*mexB* to conditions after this time point, presumably a chemically mediated plant response.

**Figure 2.**
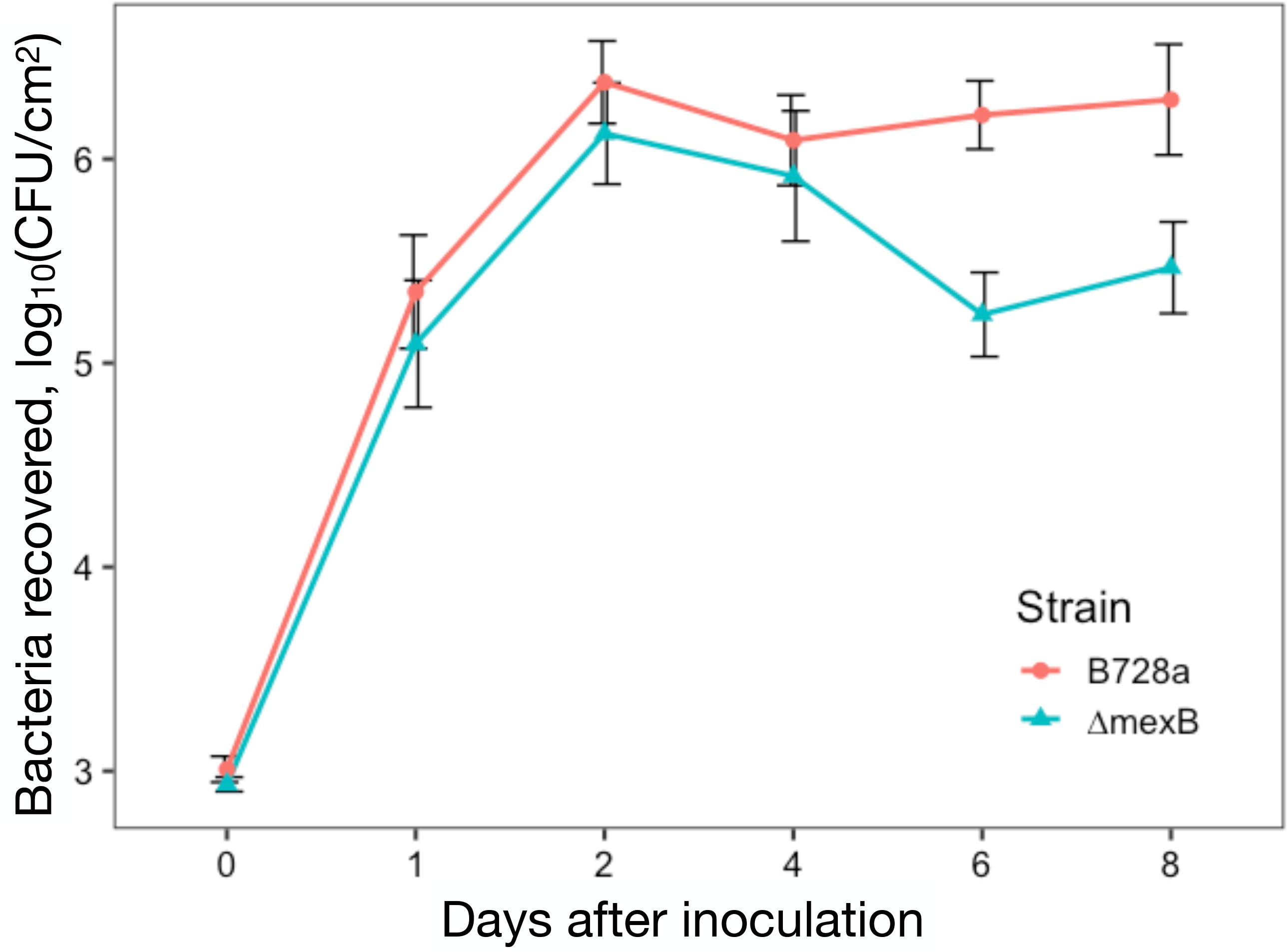
Apoplastic population sizes of *P. syringae* WT and Δ*mexB* at various times after inoculation of common bean when inoculated at 10^4^ CFU/ml. The MexAB-OprM efflux pump is not required during early stages of apoplast colonization (0 to 2 days after inoculation), and becomes more important later (ca. 6 to 8 dpi). Bacterial population sizes at 6 and 8 dpi are significantly different (Student’s t-test, p < 0.05). The vertical bars represent the standard error of the mean of log-transformed populations per cm^2^.

### Bacterial population size in leaves is driven by the number of visible lesions

When B728a is sprayed onto leaf surfaces, macroscopic lesions typically form after about seven days. Some leaves seem to be more susceptible to lesion formation than others, resulting in a rather strongly right hand skewed frequency distribution when the number of lesions is considered over a large number of leaves (Fig. S3). While it is unclear why some trifoliate leaves appear to be more susceptible to infection than others, younger leaves tended to have more lesions than more fully developed leaves. B728a Δ*mexB* formed fewer lesions per leaf than WT (Fig. S3). Inoculation with the WT strain resulted in a median of 15 and a mean of 28.8 lesions per leaf, while inoculation with the Δ*mexB* mutant resulted in a median of 2 and a mean of 5.7 lesions per leaf. It was interesting to note however, that the number of viable cells of these two strains found within discrete lesions that were excised from these leaves did not differ statistically (Fig. S4). Viable cells of these two strains could also be detected within similarly sized segments of surface sterilized asymptomatic portions of these same leaves that were sampled at the same time. While most leaf segments harvested either 7 or 21 days after inoculation harbored no detectable cells, samples from a few asymptomatic areas of the leaf contained up to 10^5^ CFU/cm^2^ (Fig. 3). Thus, while internal colonization of leaves by both strains was relatively common, it is important to note that disease symptoms were observed only when local internal population sizes in excess of about 10^5^ cells/cm^2^ were present.

**Figure 3.**
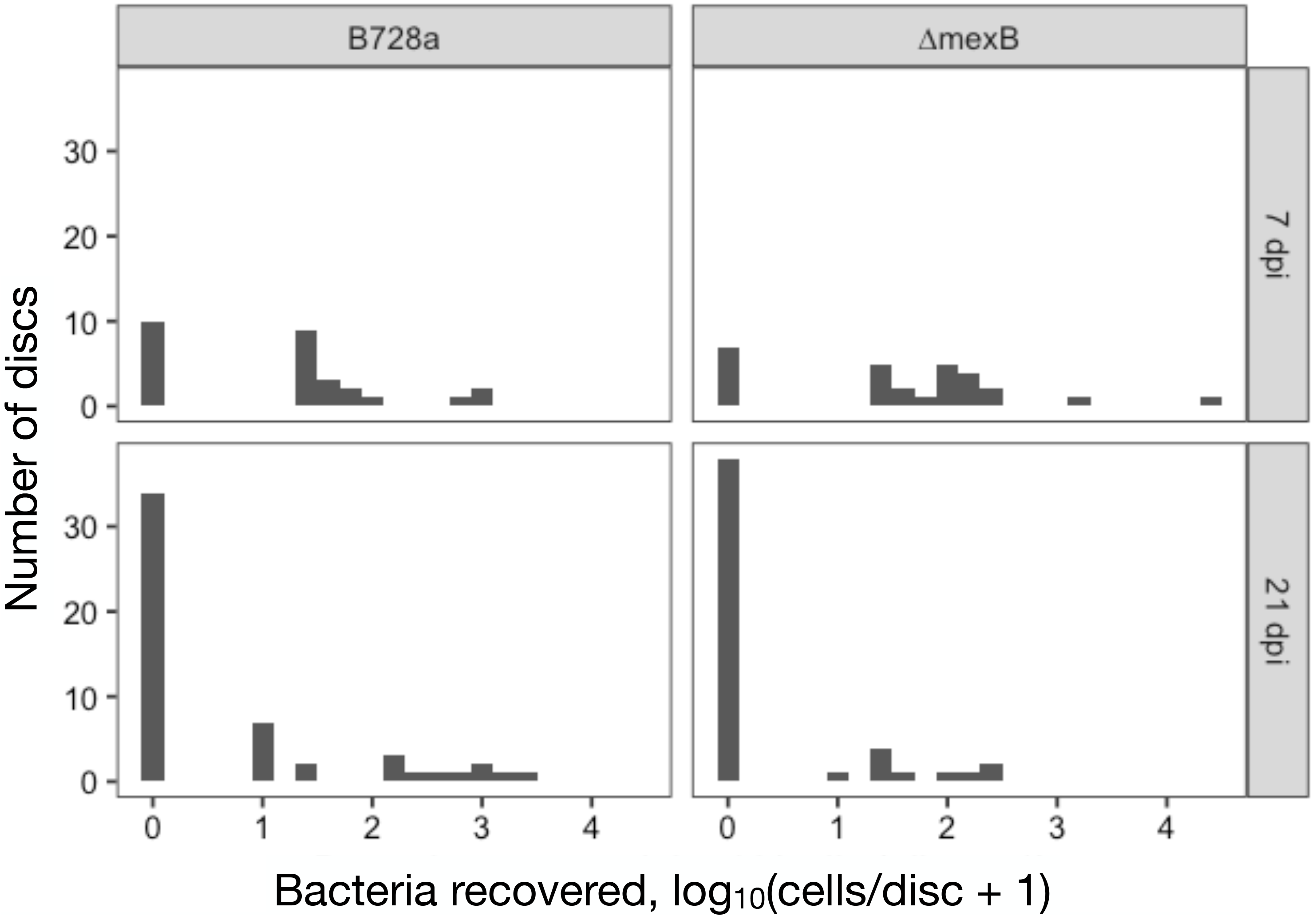
Frequency distribution of the number of bacterial cells recovered from small discs (19.6 mm^2^) excised from asymptomatic regions of common bean (*P. vulgaris*) variety Bush Blue Lake leaves that also harbored disease lesions when sampled 7 days (top panel) or 21 days (bottom panel) after topical application of *P. syringae* wild type strain B728a (left panel) or a Δ*mexB* mutant (right panel). B728a WT and Δ*mexB* cells are present within asymptomatic areas of the common bean leaf, 7 and 21 days after inoculation. N = 28 (each WT and ΔmexB) 7 dpi, 53 (WT) and 48 (Δ*mexB*) 21 dpi.

### Common bean varieties vary in their susceptibility to B728a

Given that some evidence for chemically-mediated resistance to strain B728a was observed in the susceptible *P. vulgaris* variety Bush Blue Lake 274, we explored the variation in apparent disease resistance within the *Phaseolus* genus. Wild-type strain B728a was topically applied to the leaves of 17 additional varieties of *P. vulgaris*, two varieties of *P. lunatus* (lima bean), and one variety of *P. acutifolius* (tepary bean) and the incidence of subsequent lesion formation was enumerated (Table S1). There was a wide range in the number of lesions formed per leaf among the varieties of *P. vulgaris* tested, with no lesions forming on three varieties (Pinto San Rafael, SEA5, and Tio Canela). In addition, *P. lunatus* var. Haskell and var. UC92 were more susceptible, and *P. acutifolius* var. G10 less susceptible, to lesion formation than *P. vulgaris* var. Nichols (Fig. S5).

### MexB is not required for apoplastic colonization of all plant hosts

Since varieties of common bean differed in their apparent susceptibility to lesion formation by B728a we hypothesized that their relative disease susceptibility was proportional to their ability to support the growth of this strain in the apoplast. We therefore introduced either the WT or Δ*mexB* mutant into the leaf interior of a subset of these hosts and measured increases in population size with time. Variation in the number of lesions formed on *Phaseolus* species was predicted by the extent of apoplastic growth. For example, no apoplastic growth of B728a was found in *P. vulgaris* var. Anasazi (Fig. 4A), a variety for which very few leaf lesions had been observed (Fig. S5). In contrast, *P. lunatus* supported both much higher levels of internal bacterial colonization than observed in *P. vulgaris* var. Blue Lake Bush or var. Nichols (Fig. 4B) and very high numbers of leaf lesions following spray application to leaves (Fig. S5). Interestingly, the population size of both Δ*mexB* and the WT strain in lima bean were similar and both were much higher than those on common bean varieties Blue Lake Bush and Nichols (Fig. 4B) suggesting that unlike in these other bean varieties MexB was not required for high levels of growth in lima bean. In both Blue Lake Bush and Nichols varieties the WT strain reached population levels over 10-fold greater than the Δ*mexB* mutant by 4 and 6 days after inoculation.

**Figure 4.**
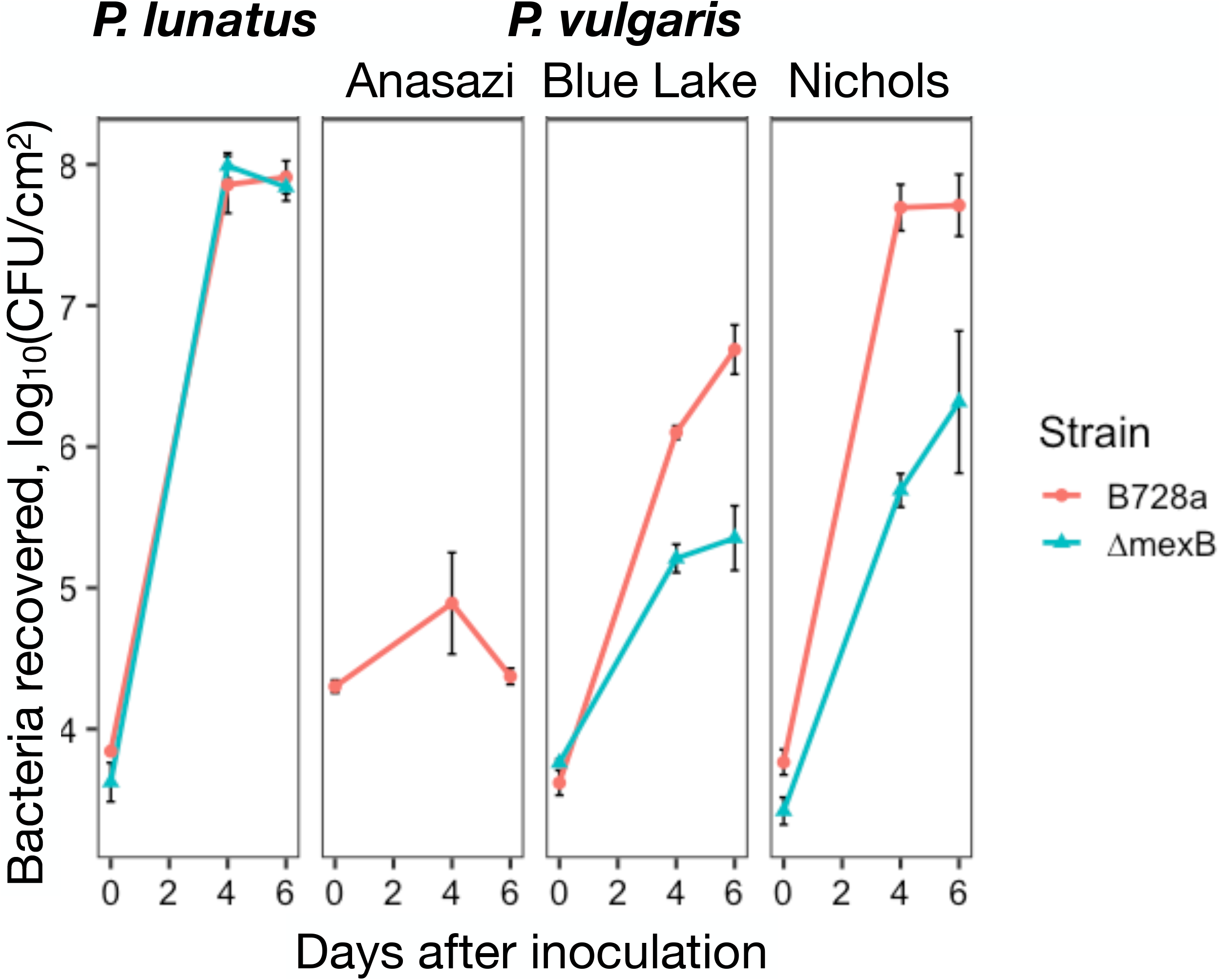
Apoplastic population size of *P. syringae* B728a WT (red) and Δ*mexB* (blue) in *P. lunatus* var. Haskell and *P. vulgaris* varieties Anasazi, Blue Lake Bush, and Nichols. The vertical bars represent the standard error of the mean of log-transformed populations per cm^2^.

To expand beyond the Fabaceae, we also tested additional plant species for their ability to support apoplastic growth of strain B728a and the requirement of MexB for such growth. *P. syringae* strain B728a was found to be capable of apoplastic growth in several additional plant hosts. While very high apoplastic population sizes were seen in both *N. benthamiana* and pepper, growth within leaves was much lower (100- to 1,000-fold increase over 6 days) in sunflower and tomato. Surprisingly, *mexB* was not required for apoplastic colonization of many of these host plants tested. While *mexB* was required for full growth in pepper, the Δ*mexB* mutant strain grew as well as the WT strain in fava bean (*V. faba*), *N. benthamiana*, sunflower (*Helianthus*), and tomato (Fig. S6). Disease symptom development was generally proportional to the extent of apoplastic growth of these strains. While extensive necrotic lesions were incited by both the WT and Δ*mexB* mutant in *N. benthamiana,* only mild discoloration was induced by the WT strain but not Δ*mexB* on pepper. No symptoms were seen on any of the other plant species that supported growth of the lower apoplastic populations of both strains. In contrast to the several other plant species for which some apoplastic growth but no symptoms were observed, very little growth of the wild type strain was seen in either black mustard (*Brassica nigra*) or red clover (*Trifolium pratense*) (Fig. S6).

### Apparent lack of chemical defenses limiting lesion formation in*N. benthamiana*

In addition to the robust apoplastic growth of strain B728a in *N. benthamiana* that continued well beyond the two to four days seen in common bean varieties (Fig. S6), we also observed greater numbers of lesions formed relative to that on common bean. Furthermore, while lesion formation on bean was relatively synchronized, with most lesions that would eventually appear on a given leaf becoming apparent by about seven days after topical application of pathogen to the leaf surface, lesions on *N. benthamiana* leaves often became apparent sooner after inoculation than those on common bean and the number of lesions on a given leaf continued to increase with time (Fig. 5). The continuing appearance of new lesions on *N. benthamiana* with increasing time after inoculation contributed to its being a much more susceptible host than common bean, and may be associated with a weak or lacking chemical defense to bacterial colonization, supported by the continued increase in apoplastic bacterial population size with time (Fig. S6) and the dispensability of MexB in the colonization of this host by strain B728a.

**Figure 5.**
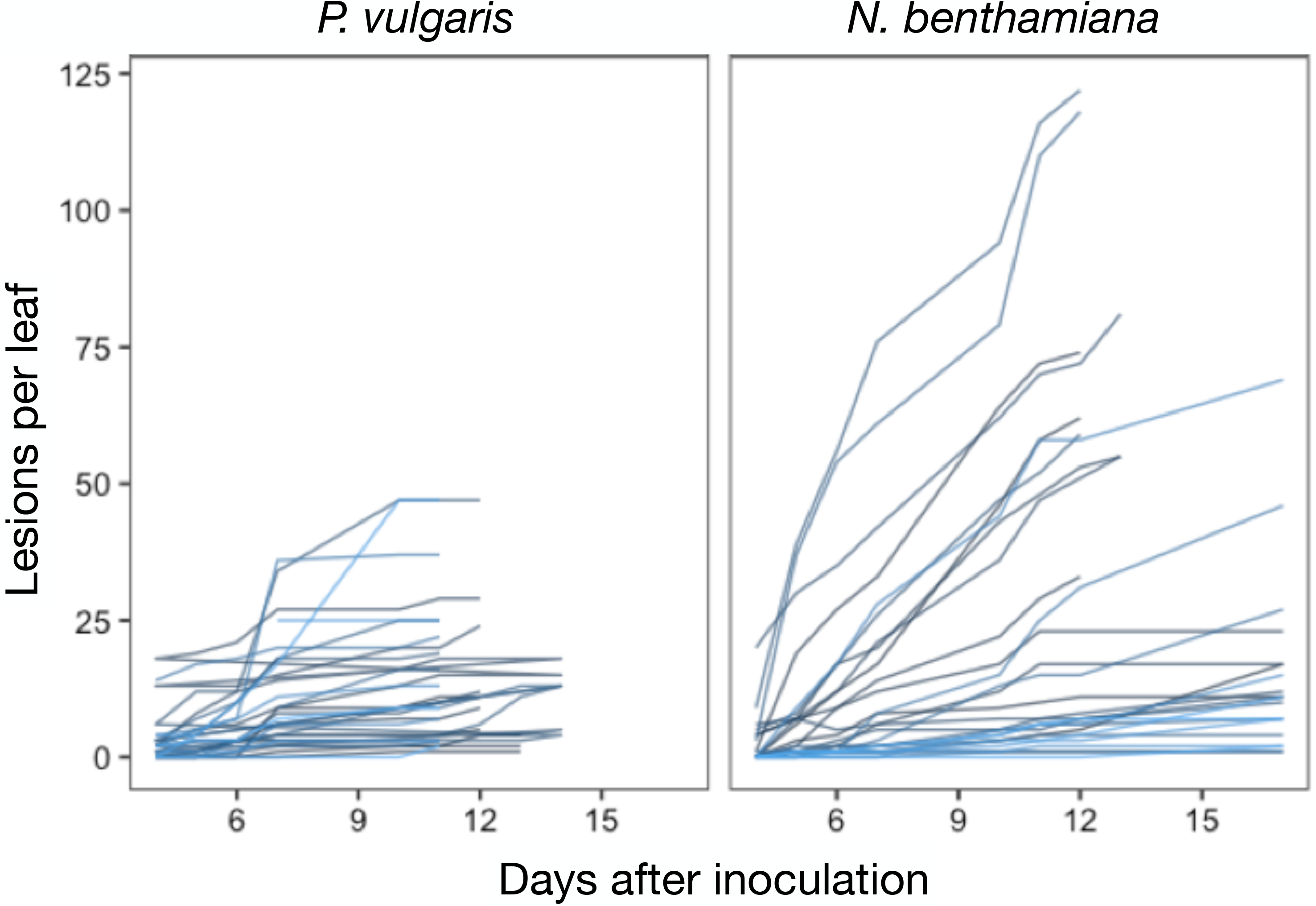
Number of lesions detected on individual leaves in common bean (*P. vulgaris*) and *N. benthamiana* when assessed at various times after inoculation. Each line represents the number of lesions on a given leaf assessed at the times after inoculation shown on the abscissa.

We hypothesized that since chemical defenses were apparently operative in common bean by at least two days after inoculation of strain B728a into leaves, bacterial populations in lesions would decline over time, as plant-produced antimicrobial compounds would accumulate with time. Furthermore, given that the Δ*mexB* mutant strain is apparently more susceptible to plant-derived toxicants than the WT strain, we hypothesized that its viability should decline at a faster rate than the WT strain. To test this, viable bacterial population sizes were determined in lesions excised from infected leaves at various times after formation. Bacterial population sizes within a given lesion of common bean, lima bean, and *N. benthamiana* generally all ranged from about 10^5^ to 10^6^ CFU soon after formation. However, the population size of both the WT and Δ*mexB* mutant strains both declined over time up to 28 days after inoculation in both common bean and lima bean, but conspicuously, neither did in *N. benthamiana* (Fig. 6).

**Figure 6.**
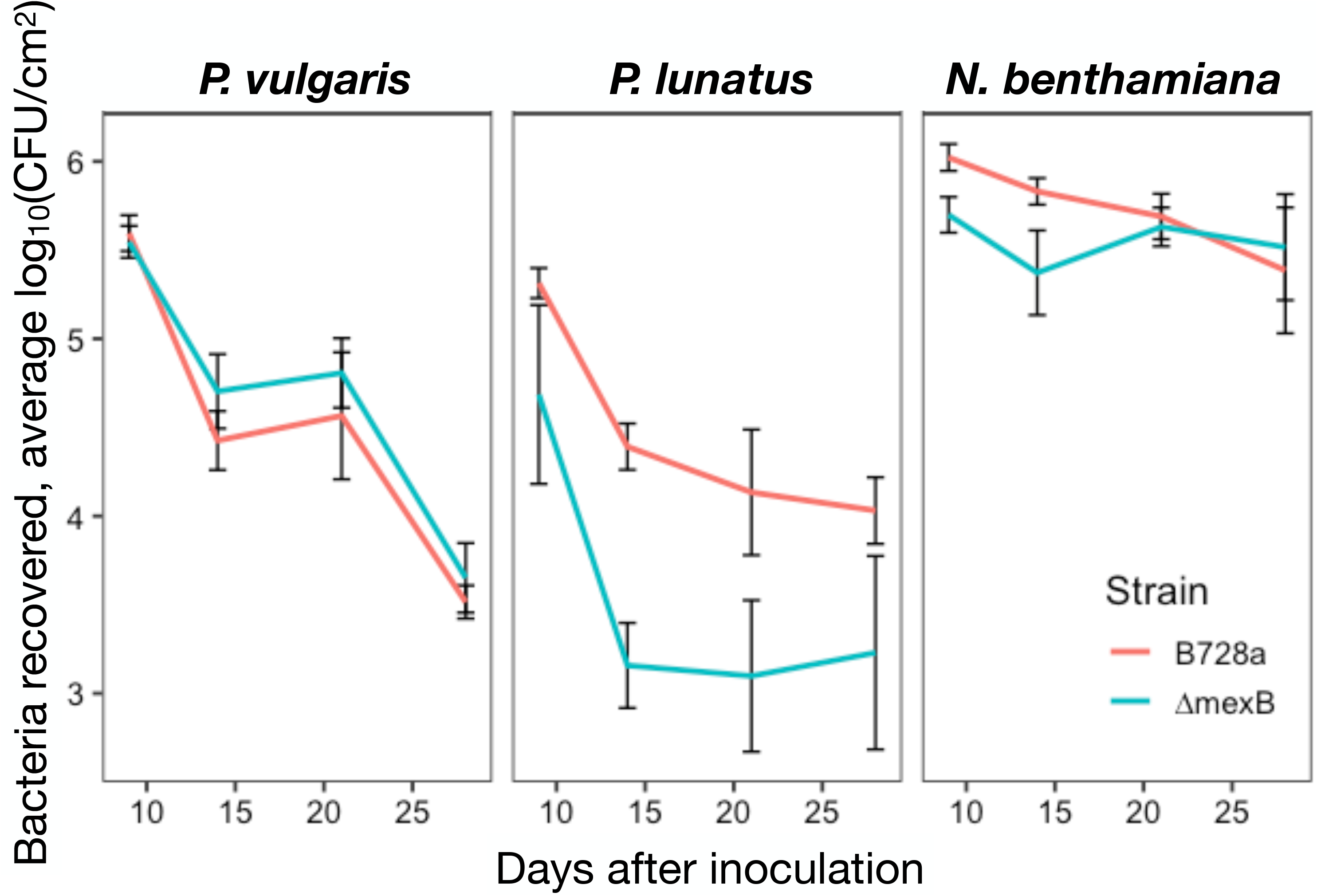
Average population size found within disease lesions formed after topical application of *P. syringae* wild type strain B728a or a Δ*mexB* mutant that were excised from leaves of common bean variety Bush Blue Lake (*P. vulgaris*), lima bean (*P. lunatus*) and *N. benthamiana* at the various times after inoculation. Bacterial populations within individual lesions decline over time in common bean and lima bean but not in *N. benthamiana*. Error bars are shown to indicate the standard error of the mean of log-transformed populations per cm^2^. Differences between populations in lesions of B728a WT and Δ*mexB* sampled at a given time were not statistically significant at any sample time, with the exception of lima bean measured at 14 days after inoculation (Welch Two Sample t-test, p = 0.002).

## Discussion

The type strain for *P. syringae* was originally isolated from diseased lilac, and most research on this species has focused on its ability to colonize diverse plant species and cause disease (Xin et al. 2018). Strain B728a was isolated from infected common bean and is a member of phylogroup 2, which consists of many pathogenic strains that are also capable of causing ice nucleation, but also includes many strains that are non-pathogenic and can be found in a variety of environmental settings not obviously associated with plants (Berge et al. 2014). The paradigm of pathogenesis for such foliar pathogens posits that infection follows the multiplication of cells on plant surfaces following their immigration from other plants or airborne or other environmental reservoirs. Only following entry of some of these epiphytic cells into the leaf interior by passage through stomata, hydathodes, or other opening such as wounds do cells gain entry into the intercellular spaces where, through their secretion of various effectors and toxins, modify the plant environment to one where extensive growth can occur. Disease symptoms seem normally to only follow the development of large internalized populations. Strains such as B728a are particularly strong epiphytic colonizers (Xin et al. 2018; Feil et al. 2005), and their high epiphytic populations can increase the likelihood of movement through stomata or leaf wounds into the apoplast and the formation of visible symptoms (Lindemann et al. 1984). Though stomata close in response to detecting microbes through the perception of pathogen-associated molecular patterns (PAMPS), *P. syringae* can secrete various strain-dependent phytotoxins to induce their reopening (Melotto et al. 2008). Foliar lesions usually follow bacterial-induced water soaking that facilitates apoplastic growth, with lesions subsequently appearing necrotic and dry (Leben 1981).

If bacterial populations above a specific threshold size were required for the formation of a visible lesion, we would expect asymptomatic areas of the leaf to host bacterial cells only below that threshold. Based on the sampling of lesions in this study, it would appear that such a localized apoplastic bacterial threshold for lesion formation is approximately 10^5^ cells per site (Fig. 3). A higher number of samples would have provided a better estimate of the threshold bacterial population size below which disease symptoms are not seen but it would not appear to be much small than that observed here. While few leaf sites (20 mm^2^) with relatively high internal populations (>10^4^ cells) were observed in this study, a surprisingly high proportion of small leaf samples (42 to 75%) harbored at least a few cells of the pathogen (Fig. 3). This suggests that the invasion of a leaf occurs at many hundreds of sites, but that growth of apoplastic populations to sizes sufficient to induce lesions is quite uncommon. As the B728a Δ*mexB* mutant formed fewer lesions on common bean in this study, and such mutants in other *P. syringae* strains also have been observed to form fewer lesions than WT strains (Stoitsova et al. 2008), we hypothesized that while this strain might enter leaves at a similar frequency as that of the WT strain, that its subsequent localized apoplastic population sizes would tend to be lower than that of the WT strain because of its apparent hyper-sensitivity to induced host defenses at such a site. As such, it would be expected to be less capable of reaching the necessary “threshold population” for symptom formation. While an attractive model, it proved difficult to test because of the relatively few sites in which substantial apoplastic growth of these strains occurred. Areas of the leaf without detectable bacterial cells likely represented areas where cells were unable to either enter or grow in the apoplast. In a low humidity environment, such as the greenhouse, bacterial populations on the leaf surface decline over time (Burch et al. 2014), and, coupled with the absence of environmental features such as large raindrops that could facilitate the entry of cells into the plant (Hirano and Upper 1983), internal colonization and subsequent lesion formation is probably less than what might be expected under certain field conditions. It is noteworthy that under these greenhouse conditions, when extrapolating across the area of the leaf as a whole, there were many more sites where *P. syringae* was detected within a leaf than the number of sites where lesions formed. It would thus appear that lesion formation might well be a relatively uncommon outcome following the introduction of this pathogen into the apoplast.

As epiphytic bacterial populations presumably must always precede lesion formation, by providing inoculum to enter and subsequently grow within the apoplast, there have been efforts to anticipate disease from knowledge of epiphytic communities. The concept of an “infection threshold” has been used to predict disease incidence at the field level, with a focus on correlating the population size of epiphytic pathogens with disease severity later in the growing season (Lindemann et al. 1984; Stromberg et al. 1999). Such predictive models are premised by knowledge of 1) the frequency with which a leaf would harbor an epiphytic population size of a given size or larger and 2) the likelihood that cells within such a population would enter the leaf, multiply, and cause lesion formation. Mechanistically, feature #2 is driven by the likelihood of lesion formation, given a particular apoplastic bacterial population. If lesion formation requires surpassing a certain internal population size threshold, then the Δ*mexB* mutant is hindered in virulence because it appears to be more sensitive to chemical defenses that it elicits during the infection process. Because of that, fewer colonization sites are likely to reach the threshold for symptom development (Fig. 7). Perhaps more importantly, the Δ*mexB* mutant also suffers from issues related to feature #1. Specifically, the Δ*mexB* mutant achieves a lower epiphytic bacterial population than the WT strain apparently because of its sensitivity to inhibitory compounds found on the leaf surface (Fig. 1). As such, fewer leaves (or leaf sites) would harbor the relatively high epiphytic populations that are associated with meaningfully high frequencies of invasion of the leaf (Fig. 7).

**Figure 7.**
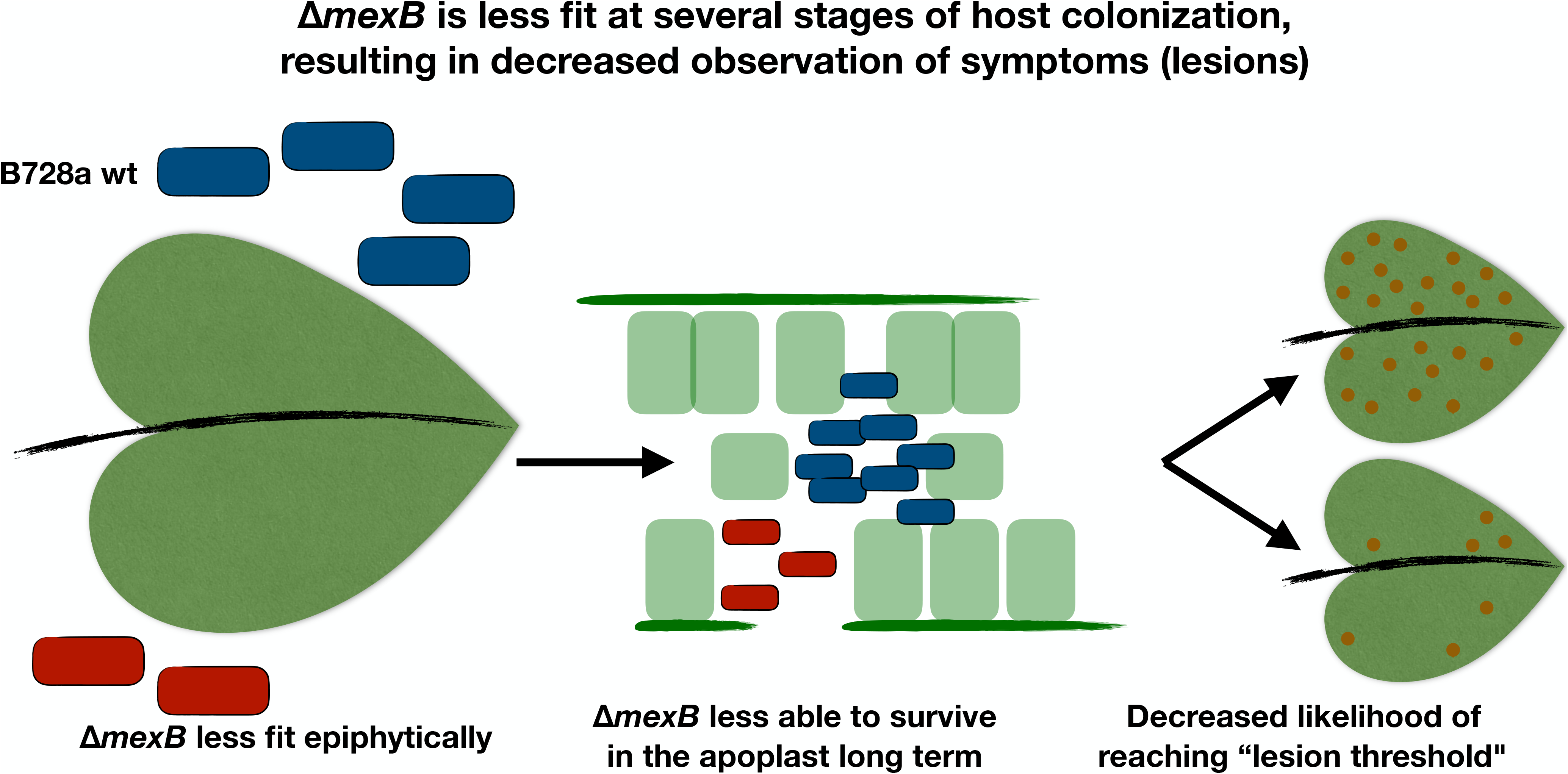
Proposed model of a population size threshold for lesion formation and the role of MexB in lesion formation. B728a Δ*mexB* is less fit both on the leaf surfaces well as in apoplastic growth, resulting in decreased incidence of lesion formation. A threshold population in the apoplast is required for visible symptoms to form on the leaf. B728a Δ*mexB* is less fit epiphytically, as well as in the apoplast (within the time frame required for lesions to form: approximately 1 week). These factors likely function synergistically, resulting in a decreased frequency of Δ*mexB* both entering the leaf and its subsequent growth in the apoplast to reach the required “lesion threshold”. These combined deficits decrease the likelihood of its epiphytic community to cause disease symptoms.

Following the observation that bacterial populations within lesions decline over time in common bean and lima bean, we hypothesized that the Δ*mexB* mutant would decline at a faster rate. This hypothesis is driven by our conjecture that the accumulation of inhibitory chemicals continues to increase with time after lesion formation, eventually becoming sufficiently high that even the WT strain cannot tolerate them. If the plant were actively reducing the bacterial load on the leaf by producing antimicrobial metabolites, then the Δ*mexB* mutant would be expected to be more susceptible, and therefore less able to survive in this local environment. Populations of Δ*mexB* within lesions were consistently lower than that of the WT strain at various times after formation in Lima bean (Fig. 6) providing some support for this conjecture. Importantly, there was a dramatically higher rate of population decline of both the WT and Δ*mexB* mutant in common bean and lima bean compared to that in *N. benthamiana* (Fig. 6). Lesions on *N. benthamiana* leaves maintained high bacterial populations for 28 days. We have generally observed that *N. benthamiana* is particularly susceptible to infection by B728a, forming many more lesions following topical inoculation. Bacterial growth within the apoplast is also more extensive and prolonged than that in either common bean or lima bean (Fig. S6). It is our conjecture that this hyper-susceptibility of *N. benthamiana* is at least partially due to a lack of a robust chemical defense. Conversely, the initial chemical defenses of common bean variety Bush Blue Lake appear to be much more robust than that of lima bean. Both *Phaseolus* species do appear to eventually accumulate antimicrobial compounds in lesions, resulting in the death of cells as lesions age. However, based on our observations, these antimicrobial compounds are not necessarily substrates of MexB in common bean. For example, plants generate redox-active molecules as part of the basal immune response (Stoitsova et al. 2008) and these may differ between plant species. Efflux transporters such as MexB appear to play an important epidemiological role in species such as *P. syringae* not only in facilitating initial colonization of plants by contributing to the tolerance of stressful chemical conditions on the leaf surface and early in lesion formation, but perhaps more importantly, by maintaining the viability of cells in lesions as they age. Such lesions provide the secondary inoculum to facilitate epidemics of diseases caused by *P. syringae*, and hence inoculum survival is an important process in the disease cycle.

The efflux transporter MexAB-OprM has been previously shown to contribute to virulence of *P. syringae* B728a on common bean leaves (Stoitsova et al. 2008). However, that study was not temporally intensive and focused only on long-term differences in bacterial populations assessed in whole infected leaves weekly for three weeks after inoculation. Stoitsova *et al.* noted that lesions formed by Δ*mexB* were less abundant but morphologically indistinguishable from those formed by the WT strain (Stoitsova et al. 2008). We confirm these findings, and we also show that bacterial populations within the lesions formed by the two strains are not significantly different. This decrease in the number of lesions, given that all lesions contain approximately the same number of bacterial cells, likely contributes to the previously-observed decrease in Δ*mexB* mutant cells per leaf (Mercier and Lindow 2000; Monier and Lindow 2004). This suggests that the difference in total population sizes in whole leaves is due primarily to the decrease in total lesions formed by Δ*mexB*. While we observed that bacterial cells were present in asymptomatic areas of the leaf, these cells represent a small fraction of the total in infected leaves when considering the high density of bacterial cells present within single lesions. Bacterial cell density on a leaf surface is spatially heterogeneous, presumably due to the heterogeneous distribution of nutrients (Mercier and Lindow 2000; Monier and Lindow 2004). The uniformity in bacterial populations within lesions suggests that there is a bacterial population threshold that must be surpassed for the formation of visible lesions on leaves, and hence the number of lesions on a leaf may be driven by the number of localized sites where epiphytic bacterial populations are sufficiently large that invasion into the apoplast is successful; lesions would form at such sites only if subsequent apoplastic growth is sufficient to exceed a threshold for symptom development. Since *mexB* is required for both full growth on the leaf surface and growth in the apoplast, it would contribute to both the frequency with which bacteria might enter the plant, and also its success in multiplying once in the plant - thereby influencing the number of lesions that would form (Fig. 7).

It is intriguing that topical inoculations of the primary leaves of common bean with *P. syringae* frequently do not yield foliar infections, unlike on trifoliate leaves. Such a finding is surprising given that these strains multiply well in primary leaves. *P. syringae* appears to grow equally well in primary and trifoliate leaves of common bean after inoculation into intercellular spaces and it is primarily due to the practicality of inoculating smaller plants having only primary leaves that studies of its colonization of this species have been typically performed at this stage of plant development. These observations suggest that disease susceptibility differs between leaf types, or leaf age. Age has been shown to impact disease susceptibility in plants. For example, *Arabidopsis* plants become more resistant to *P. syringae* as they age (Sutton and Deverall 1983). Early studies in soybean showed infection by *Sclerotinia sclerotiorum* caused symptoms on trifoliate leaves and not primary leaves (Zuiderveen et al. 2016; Balardin and Kelly 1998; Young and Kelly 1996; Guzmán et al. 1995). It is unclear whether the preferential formation of lesions on trifoliate leaves compared to primary leaves is due to easier entry into the leaf from epiphytic sources or differences in subsequent apoplastic growth, or another mechanism. Addition work is needed to confirm this phenotype in common bean, and to identify a mechanism for this variation in disease susceptibility.

Of the plant species tested here, the ability of the host to restrict growth of *P. syringae* by chemical defenses in a MexB-dependent manner was most pronounced in common bean and pepper. Furthermore, even within common bean there appears to be a substantial difference in the contribution of MexB to apoplastic growth and lesion formation (Figs. 3 and 4). This diversity in apparent disease susceptibility within both common bean and lima bean has been noted also for diseases caused by fungal pathogens such as anthracnose (*Colletotrichum lindemuthianum*) and angular leaf spot (*Phaeoisariopsis griseola*) (Hagedorn et al. 1971). Hagedorn *et al.* screened 383 lines of lima bean for tolerance to *P. syringae* and showed some variation in disease intensity in the field, but found all lines to be similarly susceptible in the greenhouse (Cichy et al. 2015). A panel of 215 common bean lines tested in the field for susceptibility to common bacterial blight (*Xanthomonas axonopodis* pv. *phaseoli*), revealed a unimodal distribution of disease susceptibility around an intermediate level (Lyon and Wood 1975). We hypothesized that the Δ*mexB* mutant was limited in growth in a host- or cultivar-specific manner due to the differential presence of antimicrobial metabolites produced by these plants. Lyon and Wood showed that *P. vulgaris* produces several phytoalexins including coumestrol and phaseollin, but that they only gradually accumulate in the apoplast within one to five days following infection by *P. phaseolicola* (Bailey and Burden 1973; Gnanamanickam and Patil 1977; Woodward 1980). These compounds can also form in bean in response to wounding, fungal infection, and viral infection (Dixon 2001). Such phytoalexins, produced after infection, therefore appear to have a primary role in limiting the success (growth) of pathogens after infection has occurred rather than preventing infection, as is the case for effector-induced resistance responses. It thus appears that there is commonly a delay in accumulation of such phytoalexins to levels that would be toxic to pathogens. It is tempting to speculate that the strong temporal dynamics of growth of *P. syringae* in plants such as common bean may be explained based on the accumulation of phytoalexins in plants only two days or more following infection. It is noteworthy that strain B728a multiplied rapidly only for about two days, at which point growth largely ceased, independent of the population size that it had achieved by that time (Figs. 2 and S2). Intriguingly, MexB only contributed to growth or survival of strain B728a in plants such as the common bean varieties Bush Blue Lake or Nichols two days or more after inoculation. The efflux transporter MexAB-OprM may be required to tolerate the presence of these plant defense compounds as they accumulate to high levels in the apoplast. As different plant species produce different defensive compounds (Tegos et al. 2002), despite its wide substrate range MexB is not capable of excluding all potential antimicrobial molecules (Helmann et al. 2019). It is tempting to speculate, therefore, that those common bean varieties and other plant species in which strain B728a achieves the lowest population size and for which MexB was required for maximal growth produce inhibitory compounds and that these compounds are also substrates for MexB. Alternatively, one or more alternative efflux transporters found in strain B728a may play a more important role than MexB in tolerating toxic compounds found in lima bean and *N. benthamiana*.

Gram-negative bacterial species are typically highly resistant to toxins *in vitro* due to the activity of a variety of conserved multidrug resistance efflux pumps. However, the chemical inhibition of these pumps can result in the increased toxicity of plant-produced antimicrobial compounds, including in *P. syringae* (Lewis and Ausubel 2006). It is possible that some of the hosts tested here, such as tomato, in which *mexB* appears dispensable, are simply producing one or more synergistically acting efflux pump inhibitors. This has been seen in barberry (*Berberis)* species, which produce the alkaloid berberine as well as the multidrug inhibitor 5’-methoxyhydnocarpin (Morris et al. 2019). In addition, we observed substantial variation in the amount of bacterial growth in the apoplast of the plants tested here, with some plant species such as *N. benthamiana* supporting more than 10,000-fold more multiplication than in species such as *V. faba* or *B. nigra* (Fig. S6). The factors limiting growth in these species remains unknown. Given that the population size of strain B728a increased 100-fold or more after inoculation in all species except *T. pretence* (Figs. S6 and S7), this suggests that they might all be considered hosts for this *P. syringae* strain. Considering the relatively low population sizes that were achieved in several of these species it is not surprising that symptoms were not seen in this study nor have they been typically seen under field conditions. These results however do add to the growing perspective that not all strains of *P. syringae*, and perhaps other plant pathogenic bacteria, are as host-specific as once thought (Morris et al. 2019).

B728a is capable of causing disease in diverse host plants (Loper and Lindow 1987). While the efflux transporter MexAB-OprM is generally considered a necessary virulence factor in this and other Gram-negative bacteria, here we show that it is contributes to virulence only in a temporally- and host-specific manner. The presence of this pump contributes to fitness on the leaf surface, but it appears that it is required in the apoplast only once host chemical defenses have been activated. Further exploration is necessary to better understand the specific metabolic differences between these hosts, the substrate specificities of MexB *in planta*, as well as whether other pumps are complementary to, or replace, the role of MexAB-OprM for tolerating the toxins present in other settings.

## Materials and Methods

### Bacterial strains and growth media

*Pseudomonas syringae* pv. *syringae* B728a was originally isolated from a common bean leaf (*Phaseolus vulgaris*) in Wisconsin (King et al. 1954). B728a and the kanamycin-resistant derivative mutant strain Δ*mexB* were grown on King’s B (KB) agar or in broth (Hockett et al. 2013), at 28°C. *Escherichia coli* strains S17-1, and TOP10 were grown on LB agar or in LB broth at 37°C. When appropriate, the following antibiotics were used at the indicated concentrations: 100 μg/ml rifampicin, 50 μg/ml kanamycin, 15 μg/ml tetracycline, 40 μg/ml nitrofurantoin, and 21.6 μg/ml natamycin (an anti-fungal).

### Construction of the B728a Δ*mexB* mutant strain

The construction of the Δ*mexB* mutant strain followed an overlap extension PCR protocol as described previously (Datsenko and Wanner 2000). DNA fragments upstream (1 kb) and downstream (1.3 kb) of *Psyr_4008* (*mexB*) were amplified along with a kanamycin resistance cassette from pKD13 (Chen et al. 2010). These three fragments were joined by overlap extension PCR. The resulting fragment was blunt-end ligated into the SmaI site of pT*sacB* (Simon et al. 1983), and transformed into the *E. coli* subcloning strain TOP10, and then the *E. coli* conjugation donor strain S17-1. This suicide plasmid was conjugated into B728a on KB overnight, and then selected by growing the exconjugants for three days on KB containing kanamycin and nitrofurantoin (*E. coli* counter selection). Putative double-crossover colonies that were kanamycin resistant and tetracycline sensitive were selected for screening using external primers and further confirmed by PCR and Sanger sequencing. Strains, plasmids, and primers used are listed in Table S2.

### Bacterial *in vitro* growth measurements

B728a WT and Δ*mexB* overnight cultures were grown in KB containing rifampicin and standardized to OD_600_ = 0.3. 20 μl was inoculated into a 96-well plate containing 100 μl King’s B broth (5 replicate wells each). Cells were grown at 28°C with shaking, with absorbance measurements made at 600 nm every 30 minutes. Absorbance is reported as the mean and standard deviation of absorbance at 600 nm with the average absorbance of two KB blanks subtracted from the total absorbance for each sample.

### Plant growth conditions

All *Phaseolus* varieties with the exception of Bush Blue Lake were acquired as seed from UC Davis. Common bean (*P. vulgaris*), lima bean (*P. lunatus*), tepary bean (*P. acutifolius*), fava bean (*V. faba*), sunflower (*Helianthus*), mustard (*B. nigra*), and clover (*T. pratense*) seeds (5 - 7 per 10 cm diameter pot) were planted in Super Soil (Scotts MiracleGro) and grown in a greenhouse until primary leaves were fully expanded before inoculation (approximately two weeks for bean species). *N. benthamiana,* pepper (*C. annuum* cv. Cal Wonder), and tomato (*S. lycopersicum* cv. Moneymaker) seeds were sown on Sunshine Mix #4 (SunGro Horticulture) to germinate, and then transplanted into 10 cm diameter pots containing Super Soil. *P. vulgaris* var. Blue Lake Bush 274 was used for all experiments in common bean unless otherwise specified. Leaves were kept dry to minimize epiphytic contamination. We used 1000 W metal halide lights to provide supplemental lighting for a 16-hour day length. Greenhouse temperatures ranged from 18°C at night to approximately 40°C during the day.

### Spray inoculations to measure bacterial epiphytic population size

Strains were grown overnight on KB plates containing rifampicin, washed in 10 mM KPO_4_, (pH 7.0) and resuspended in 10 mM KPO_4_. Cultures were diluted to a concentration of 10^6^ CFU/ml (OD_600_ = 0.001, by dilution from OD_600_ = 0.1) and sprayed onto the surface of primary and trifoliate leaves until runoff. Plants were placed in a high humidity mist chamber for two days. After this time, leaves were removed and placed into glass tubes containing 25 mL 10 mM KPO_4_, where cells were dislodged from the leaves using a water bath sonicator (Branson 5510, output frequency 40 kHz) for 15 min. Bacterial populations were measured by dilution plating onto KB agar containing rifampicin and natamycin.

### Bacterial apoplastic growth measurements

Strains were grown overnight on KB plates containing rifampicin, washed in 10 mM KPO_4_, and standardized to 2×10^5^ CFU/ml (OD_600_ = 0.0001, by dilution from OD_600_ = 0.1) (*Phaseolus* spp. and *Helianthus*), 10^4^ CFU/ml (*C. annuum*, *S. lycopersicum*, and *V. faba*), or 10^3^ CFU/ml (*N. benthamiana*, *Brassica*, and *Trifolium*) in 1 mM KPO_4_. Cells were inoculated into leaves using a blunt syringe. Leaf samples (3 discs per leaf) were excised using a 5 mm-diameter cork borer into tubes containing 200 μl 10 mM KPO_4_ and two 3 mm glass beads, and macerated for 30 seconds at 2400 rpm in a Mini-Beadbeater-96 (Biospec Products) before dilution plating on KB with rifampicin and natamycin.

### Spray inoculations to measure lesion formation and bacterial populations within symptomatic and asymptomatic areas of leaf tissue

Bacterial strains were grown overnight on KB plates containing rifampicin, washed in 10 mM KPO_4_, and resuspended in 10 mM KPO_4_. Cultures were diluted to a concentration of 10^7^ CFU/ml in 10 mM KPO_4_ (OD_600_ = 0.01, by dilution from OD_600_ = 0.1) and sprayed onto the surface of trifoliate leaves of Bush Blue Lake bean until runoff. Due to its higher susceptibility, *N. benthamiana* plants were sprayed with inoculum diluted to 10^5^ CFU/ml. Plants were placed in a high humidity mist chamber for two days, and then moved to the greenhouse. Individual lesions and asymptomatic areas of leaves were sampled at least 7 days after spraying, the specific timing noted for each experiment. Leaves were surface sterilized by exposure to UV irradiation in a biosafety cabinet for 30 seconds per side as in other studies (Wilson et al. 1999). Leaf samples (1 disc = 1 lesion) were excised with a 5 mm-diameter cork borer and placed in tubes containing 200 μl 10 mM KPO_4_ and two 3 mm glass beads, and ground for 30 seconds at 2400 rpm in a Mini-Beadbeater-96 before dilution plating on KB with rifampicin and natamycin.

## Acknowledgements

We thank Dr. Paul Gepts at UC Davis for providing *Phaseolus* seeds. Plants were grown and maintained with the assistance of the UC Berkeley greenhouse staff, led by Tina Wistrom. We thank Kendra Paskvan and Caitlin Ongsarte for assistance with plant inoculations and sampling. Funding for TCH was partially provided by the Arnon Graduate Fellowship and the William Carroll Smith Fellowship.

