## Supplementary Information for "Differential virulence contributions of the efflux transporter MexAB-OprM in *Pseudomonas syringae* infecting a variety of host plants"

Tyler C. Helmann, 0000-0002-8431-6461

Dana M. King, 0000-0003-0853-1511

Steven E. Lindow, 0000-0001-8333-6674

**Supplementary data and figures**

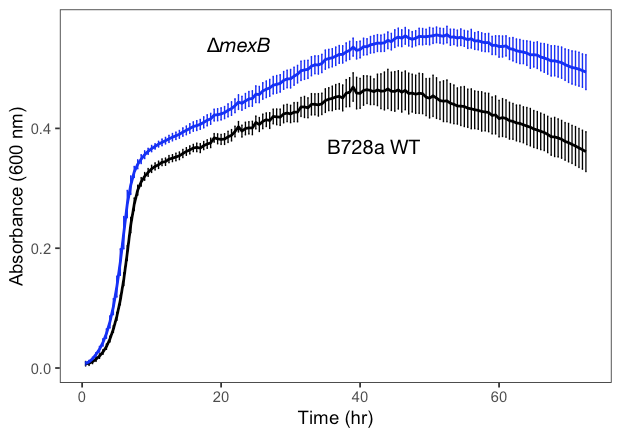

Figure S1. B728a ∆*mexB* (blue) grows as well as WT (black) in King’s B (KB) broth. Average absorbance at 600 nm is shown here for 5 replicate samples each, with the average absorbance of two KB blanks subtracted from the total absorbance for each sample. Vertical lines indicate the standard deviation.

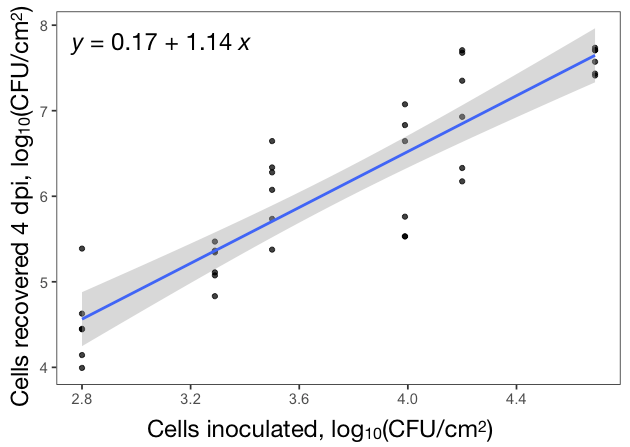

Figure S2. Population size of WT *P. syringae* strain B728a achieved after growth in leaves of common bean (*P. vulgaris*) variety Bush Blue Lake for 4 days at various concentrations is directly proportional with the concentration of inoculum introduced into the plant. A linear model was used to create the regression line. Pearson’s correlation coefficient = 0.90, p = 1.2 X 10^-13^.

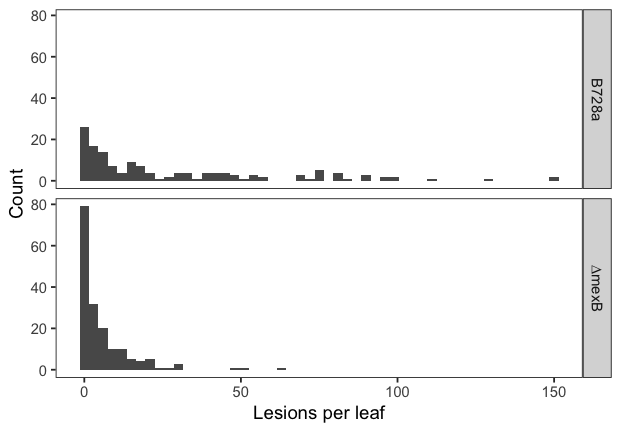

Figure S3. Frequency distribution of the number of lesions formed on individual leaves of common bean (*P. vulgaris*) variety Bush Blue Lake after topical application of *P. syringae* WT strain B728a (top panel) or a ∆*mexB* mutant (bottom panel). B728a ∆*mexB* forms fewer lesions than WT on common bean. The number of lesions formed by B728a WT and ∆*mexB* are significantly different (Wilcoxon rank sum test with continuity correction, p = 7*10^-15^). N = 146 leaves (WT) and 173 leaves (∆*mexB*), measured 10 days after inoculation.

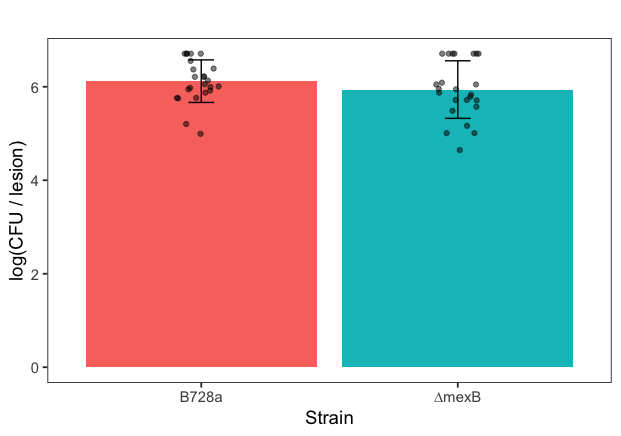

Figure S4. Population size of cells found within newly formed disease lesions formed after topical application of *P. syringae* wild type strain B728a or a ∆*mexB* mutant that were excised in 19.6 mm^2^ discs from leaves of common bean variety Bush Blue Lake. Bacterial populations within lesions sampled 7 days after inoculation are not statistically different between B728a WT and ∆*mexB* (Welch Two Sample t-test, p = 0.26). Error bars indicate the standard deviation of log-transformed CFU per lesion. Black dots represent the population size in a given lesion. N = 24 individual lesions per strain.

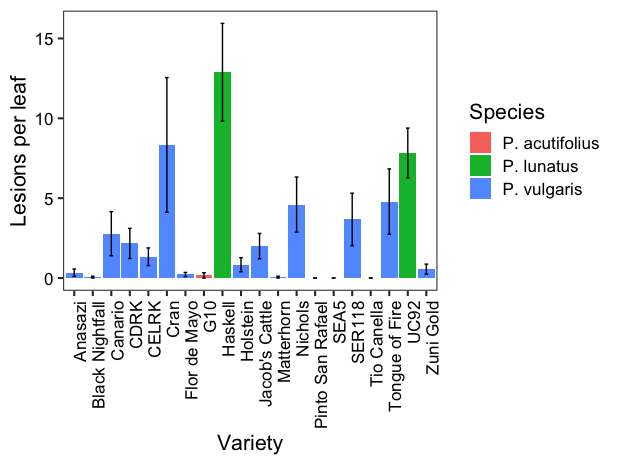

Figure S5. Number of lesions formed by *P. syringae* wild type strain B728a on leaves of varieties of three *Phaseolus* species, measured 10 days following a spray inoculation. The number of leaves assessed ranged from 6 to 18 per variety. Error bars indicate standard error of the mean.
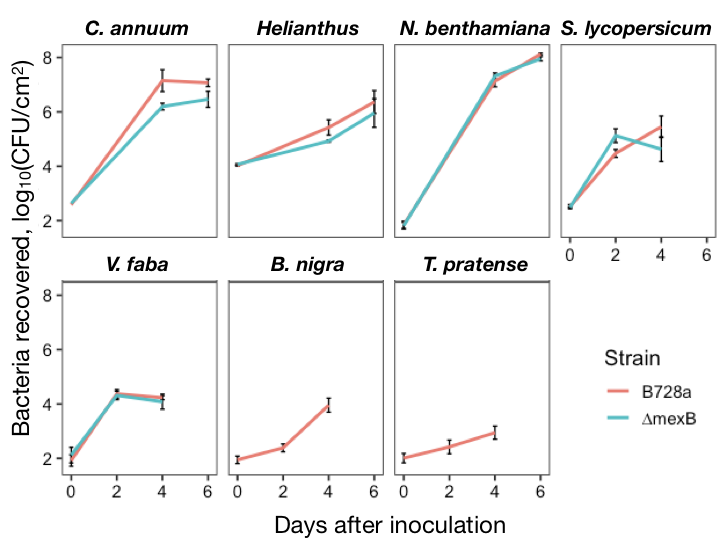

Figure S6. Apoplastic population size of *P. syringae* wild type strain B728a (red) and ∆*mexB* (blue) recovered from a variety of plant species at various times after inoculation. MexB is required for full apoplastic growth in pepper (*C. annuum*), but not fava bean (*V. faba*), *N. benthamiana*, sunflower (*Helianthus*), or tomato (*S. lycopersicum*). Total bacterial growth was relatively low (100- to 1,000-fold increase over 6 days) in sunflower and tomato. Strain B728a exhibited modest growth in the apoplast of mustard (*B. nigra*) and almost no growth in clover (*T. pratense*). The vertical bars represent the standard error of the mean of log-transformed populations per cm^2^.

Table S1. *Phaseolus* varieties used in this study.

| Species | Name | Source |
| --- | --- | --- |
| *P. vulgaris* | Blue Lake Bush 274 | Trinidad Benham Corp. |
| *P. vulgaris* | Anasazi | Paul Gepts |
| *P. vulgaris* | Black Nightfall | Paul Gepts |
| *P. vulgaris* | UCD 707 Canario | Paul Gepts |
| *P. vulgaris* | California Early Light Red Kidney (CELRK) | Paul Gepts |
| *P. vulgaris* | California Dark Red Kidney (CDRK) | Paul Gepts |
| *P. vulgaris* | UCD 0801 Cran | Paul Gepts |
| *P. vulgaris* | Flor de Mayo de Euglina | Paul Gepts |
| *P. vulgaris* | UCD Holstein | Paul Gepts |
| *P. vulgaris* | UCD Jacob's Cattle 0908 | Paul Gepts |
| *P. vulgaris* | Matterhorn | Paul Gepts |
| *P. vulgaris* | UC Nichols | Paul Gepts |
| *P. vulgaris* | Pinto San Rafael | Paul Gepts |
| *P. vulgaris* | SEA 5 | Paul Gepts |
| *P. vulgaris* | SER 118 | Paul Gepts |
| *P. vulgaris* | Tio Canela 75 | Paul Gepts |
| *P. vulgaris* | Tongue of Fire | Paul Gepts |
| *P. vulgaris* | Zuni Gold | Paul Gepts |
| *P. lunatus* | UC Haskell | Paul Gepts |
| *P. lunatus* | UC 92 | Paul Gepts |
| *P. acutifolius* | G40010 | Paul Gepts |

Table S2. Strains, plasmids, and primers used in this study.

| Strains | | |
| --- | --- | --- |
| Organism | Description | Source |
| *E. coli* TOP10 | For general cloning | Invitrogen |
| *E. coli* S17-1 | Conjugation donor strain | (Loper and Lindow 1987) |
| *P. syringae* B728a | Wild type strain (Rif^R^) | (Datsenko and Wanner 2000) |
| *P. syringae* B728a | ∆*mexB* (Rif^R^ Kan^R^) | This work |

| Plasmids | | | |
| --- | --- | --- | --- |
| Name | Description | Antibiotic | Source |
| pKD13 | Source of kanamycin resistance | Kan | (Chen et al. 2010) |
| pT*sacB* | Suicide plasmid to introduce DNA into *P. syringae* | Tet | (Chen et al. 2010) |
| pT:4008-kan | To delete *Psyr_4008*, contains *Psyr_4008* flanking regions bordering kan^R^ cassette, inserted into SmaI site of pT*sacB* | Tet Kan | This work |

| Primers | Sequence |
| --- | --- |
| FRT-KanF | GTGTAGGCTGGAGCTGCTTC |
| FRT-KanR | ATTCCGGGGATCCGTCGACC |
| 4008 up F | AGCACAGCCCTATACCCTGA |
| FRT 4008 up R | **GAAGCAGCTCCAGCCTACAC**TTACTCCCCTTTGCTGCCTG |
| FRT 4008 down F | **GGTCGACGGATCCCCGGAAT**TGCAGTTACCGCGTTCATTC |
| 4008 down R | AGGAAGGTCAGGTTGCTGTC |
| 4008 check F | CAGTGGCTGTAGCAAGAAGGA |
| 4008 check R | TTTGTTGCGCACTGAACAGC |

Bold sequence complements FRT-Kan sequence for SOE protocol.
